# Cross-Cancer Evaluation of Polygenic Risk Scores for 17 Cancer Types in Two Large Cohorts

**DOI:** 10.1101/2020.01.18.911578

**Authors:** Rebecca E. Graff, Taylor B. Cavazos, Khanh K. Thai, Linda Kachuri, Sara R. Rashkin, Joshua D. Hoffman, Stacey E. Alexeeff, Maruta Blatchins, Travis J. Meyers, Lancelote Leong, Caroline G. Tai, Nima C. Emami, Douglas A. Corley, Lawrence H. Kushi, Elad Ziv, Stephen K. Van Den Eeden, Eric Jorgenson, Thomas J. Hoffmann, Laurel A. Habel, John S. Witte, Lori C. Sakoda

## Abstract

Genetic factors that influence etiologic mechanisms shared across cancers could affect the risk of multiple cancer types. We investigated polygenic risk score (PRS)-specific pleiotropy across 17 cancers in two large population-based cohorts. The study population included European ancestry individuals from the Genetic Epidemiology Research on Adult Health and Aging cohort (16,012 cases, 50,552 controls) and the UK Biobank (48,969 cases, 359,802 controls). We selected known independent risk variants from published GWAS to construct a PRS for each cancer type. Within cohorts, each PRS was evaluated in multivariable logistic regression models with respect to the cancer for which it was developed and each other cancer type. Results were then meta-analyzed across cohorts. In the UK Biobank, each PRS was additionally evaluated relative to 20 cancer risk factors or biomarkers. All PRS replicated associations with their corresponding cancers (*p*<0.05). Eleven cross-cancer associations – ten positive and one inverse – were found after correction for multiple testing (p<0.05/17=0.0029). Two cancer pairs showed bidirectional associations; the melanoma PRS was positively associated with oral cavity/pharyngeal cancer and vice versa, whereas the lung cancer PRS was positively associated with oral cavity/pharyngeal cancer, and the oral cavity/pharyngeal cancer PRS was inversely associated with lung cancer. We identified 65 associations between a cancer PRS and non-cancer phenotype. In this study examining cross-cancer PRS associations in two cohorts unselected for phenotype, we validated known and uncovered novel patterns of pleiotropy. Our results have the potential to inform investigations of risk prediction, shared etiology, and precision cancer prevention strategies.

**STATEMENT OF SIGNIFICANCE:** By examining cross-cancer polygenic risk score associations, we validated known and uncovered novel patterns of pleiotropy. Our results may inform investigations of risk prediction, shared etiology, and precision prevention strategies.

## INTRODUCTION

Neoplasms are remarkably diverse in their clinical presentation, but they share biological hallmarks acquired during the transformation of normal cells into neoplastic ones (1). Inherited genetic factors underpinning shared hallmarks could alter cancer risk in a pleiotropic manner. Indeed, genome-wide association studies (GWAS) of individual cancer types have identified loci associated with other cancer types, including 5p15 (*TERT-CLPTM1L)* (2), 6p21 (HLA complex) (3,4), and 8q24 (5). Non-GWAS approaches have yielded further pleiotropic cancer risk variants, and genetic correlation studies have identified cancer pairs with shared heritability (6-8).

Polygenic risk scores (PRS) capture a different aspect of pleiotropy. By combining variants into scores that summarize genetic susceptibility, PRS typically explain a larger proportion of disease risk than single low-penetrance variants. Relative to genetic correlations, PRS offer greater specificity by selecting a refined set of disease-specific risk variants. PRS analyses therefore have the potential to inform etiology and identify possible precision prevention targets shared across cancers. They are also plausibly valuable for risk prediction; there is potential clinical advantage in knowing that an individual with a high PRS for one cancer is at risk for another. While PRS have been extensively investigated for individual cancers, cross-cancer portability of PRS has not been well studied.

To comprehensively investigate pleiotropy across cancers, we leveraged results from 273 published GWAS to systematically construct PRS specific to 17 cancer types. We then evaluated associations between each PRS and the risk of each cancer type in European ancestry individuals from two large independent cohorts with genome-wide array data – the Genetic Epidemiology Research on Adult Health and Aging (GERA) cohort and the UK Biobank. We also assessed associations between each genetic variant contributing to a PRS and the risk of each cancer type and characterized pleiotropy between each PRS and 20 cancer risk factors or biomarkers.

## MATERIALS AND METHODS

### Study Populations

GERA is a prospective cohort of 102,979 adults drawn from >400,000 Kaiser Permanente Northern California (KPNC) health plan members who participated in the Research Program on Genes, Environment and Health. Participants answered a baseline survey regarding lifestyle and medical history, provided a saliva specimen between 2008 and 2011, and were successfully genotyped (9,10). Following quality control (QC; described below), the GERA analytic population included 16,012 cases and 50,552 controls.

The UK Biobank is a population-based prospective cohort of 502,611 individuals from the United Kingdom, ages 40 to 69 at recruitment between 2006 and 2010 (11). Participants were evaluated at baseline visits during which assessment center staff introduced a touch-screen questionnaire, conducted a brief interview, gathered physical measurements, and collected biological samples. Following QC, the UK Biobank analytic population included 48,969 cases and 359,802 controls.

This study was approved by the KPNC and University of California Institutional Review Boards and the UK Biobank data access committee.

### Phenotyping

GERA cancer cases were identified using the KPNC Cancer Registry. Following Surveillance, Epidemiology, and End Results Program (SEER) standards, the KPNC Cancer Registry contains data on all primary cancers (i.e., diagnoses that are not secondary metastases of other cancer sites; excluding non-melanoma skin cancer) diagnosed or treated at any KPNC facility since 1988. In this study, we captured all diagnoses recorded through June 2016. Cancer cases in the UK Biobank were identified via linkage to various national cancer registries (11). Data in the registries are compiled from hospitals, nursing homes, general practices, and death certificates, among other sources. Diagnoses go as far back as the early 1970s, and the latest cancer diagnosis in our data from the UK Biobank occurred in August 2015.

In both cohorts, individuals with at least one recorded prevalent or incident diagnosis of a borderline, in situ, or malignant primary cancer were defined as cases. To align with GERA, we converted all UK Biobank diagnoses described by International Classification of Diseases (ICD)-9 or ICD-10 codes into ICD-O-3 codes. We then classified cancers in both cohorts by organ site according to the SEER site recode paradigm. Because second and subsequent cancers could have been miscoded metastases of a first cancer or a direct result of prior cancer treatment, we evaluated only the first primary cancer diagnosed for each individual. The analyses did, however, include 23 GERA participants and 64 UK Biobank participants who had two primary cancers diagnosed on the same date. To ensure sufficient statistical power, we grouped all oral cavity and pharyngeal cancers and, separately, all esophageal and stomach cancers into single site codes. Overall, our analyses included 17 of the most common site codes (excluding non-melanoma skin cancer). Data on testicular cancer cases were obtained from the UK Biobank only due to the small number of cases in GERA.

Controls were restricted to individuals who had no record of cancer in any of the relevant registries, who did not self-report a prior history of cancer (other than non-melanoma skin cancer) by survey, and, if deceased, who did not have cancer listed as a cause of death. For analyses of sex-specific cancer outcomes (breast, cervix, endometrium, ovary, prostate, and testis), controls were restricted to individuals of the relevant sex.

In the UK Biobank, we examined PRS associations with anthropometric traits, physical measures, self-reported health-related behaviors, and serum biomarkers. Physical assessments yielded measures of height (Field ID: 50.0), body mass index (BMI; Field ID: 21001.0), waist to hip ratio (Field ID: 48.0 divided by Field ID: 49.0), diastolic blood pressure (Field ID: 4079.0), and systolic blood pressure (Field ID: 4080.0). Body fat percentage (Field ID: 23104.0) was quantified with whole-body bio-impedance measures using the Tanita BC418MA body composition analyzer. Self-reported data on cigarette smoking and alcohol consumption were used to derive variables for smoking status (ever/never), cigarettes per day, and weekly alcohol intake (grams). We additionally evaluated eight serum biomarkers, as measured according to protocols that have been previously described (12) – C-reactive protein (CRP; mg/L), high-density lipoproteins (HDL; mmol/L), low-density lipoproteins (LDL; mmol/L), glycated hemoglobin (HbA1c; mmol/mol), insulin-like growth factor-1 (IGF-1; nmol/L), sex hormone binding globulin (nmol/L), testosterone (nmol/L) in men, and testosterone (nmol/L) in women. All biomarker analyses were restricted to samples from the first aliquot, as these samples were least affected by unintended sample dilution issues (12). We excluded values outside the bioanalyzer reportable range, as well as measures that required additional analytic correction due to sample handling or processing issues. CRP, HbA1c, and IGF-1 were log-transformed to achieve a normal distribution.

### Genotyping and Imputation

For GERA, genotyping was performed using one of four Affymetrix Axiom arrays (Affymetrix, Santa Clara, CA, USA) optimized for individuals of European, African, East Asian, and Latino race/ethnicity. Details about the array design, estimated genome-wide coverage, and QC procedures have been published previously (10,12-14). Variants that were not directly genotyped (or excluded by QC procedures) were imputed to generate genotypic probability estimates. After pre-phasing genotypes with SHAPE-IT v2.5 (15), IMPUTE2 v2.3.1 was used to impute variants relative to the cosmopolitan reference panel from the 1000 Genomes Project (phase I integrated release; http://1000genomes.org/) (16). Ancestry principal components (PCs) were computed using Eigenstrat v4.2, as previously described (9,17).

For the UK Biobank, genotyping was conducted for 436,839 individuals with the UK Biobank Axiom array and for 49,747 individuals with the UK BiLEVE array (11). The former is an updated version of the latter, such that the two arrays share over 95% of their marker content. UK Biobank investigators undertook a rigorous QC protocol (11). Imputation was performed primarily based on the Haplotype Reference Consortium reference panel, and the merged UK10K and 1000 Genomes Project (phase 3) reference panels were used for secondary data (11). Ancestry PCs were computed using *fastPCA* (18) based on a set of 407,219 unrelated samples and 147,604 genetic markers (11).

### Quality Control

Additional QC procedures included restricting to self-reported European ancestry individuals with matching self-reported and genetic sex. To further minimize population stratification, we excluded individuals for whom either of the first two ancestry PCs fell >5 standard deviations outside of the mean. We also removed samples with call rates <97%, heterozygosity >5 standard deviations from the mean, and/or first-degree relatives in the datasets.

### Variant Selection for PRS

PRS were constructed based on variants associated with each cancer type in existing published GWAS. To identify relevant GWAS, we began by searching the National Human Genome Research Institute-European Bioinformatics Institute Catalog of published GWAS (19). For every GWAS of a cancer of interest (or one of its sub-phenotypes; e.g., *poorly differentiated* prostate cancer) that discovered at least one genome-wide significant (*p*≤5×10^−8^) risk variant, we reviewed both the original primary manuscript and supplementary materials. We then identified additional relevant GWAS by 1) reviewing the reference section of each article, and 2) searching PubMed to find other studies in which each article had been cited (**Supplementary Table 1**). Only one out of 273 studies identified included data that overlapped with ours; UK Biobank data accounted for 21% of the Huyghe, *et al*. study population and was only used in the second stage of their GWAS (20).

After abstracting genome-wide significant variants from all studies published by June 30, 2018, we reduced the file to include one log-additive association per combination of variant identifier, phenotype / sub-phenotype, and ancestry group (**Supplementary Figure 1**). For associations reported in more than one study of the same ancestry, we selected the one with a known risk allele and effect estimate with the smallest p-value.

We retained only autosomal variants identified in populations of at least 70% European ancestry. We then excluded 2,979 associations for which the source literature did not report an effect estimate and/or for which an effect allele could not be determined. For the remaining 13,827 associations, we assessed variant availability in both the GERA and UK Biobank genotypic data. For lead variants that could not be identified by variant identifier or position, we used LDlink (21) and HaploReg (22) to identify proxy variants with r^2^≥0.8. From original or proxy variants available in GERA and UK Biobank, we excluded any not in a 1000 Genomes reference population or with minor allele frequencies (MAF) that differed by >0.10, and we further restricted to biallelic risk variants with MAF ≥0.01. In a last step prior to linkage disequilibrium (LD) pruning, we excluded A/T and C/G variants with MAF ≥0.45 – due to strand flips, the appropriate effect alleles in our data could not be determined.

We used PriorityPruner (23) and LDlink (21) to select a set of independent risk variants with LD <0.3 for each cancer type. The process preferentially selected variants with the smallest *p*-values and highest imputation scores associated with the broadest phenotype (e.g., overall prostate cancer over poorly differentiated prostate cancer).

### Statistical Analysis

For each cancer type, we calculated the PRS based on additive dosages of the individual risk variants: ∑(# risk alleles*log(odds ratios [OR]) from the literature) for i = 1 to n risk alleles. Each PRS was then standardized based on its mean and standard deviation, and evaluated in multivariable logistic regression models with respect to the cancer for which it was developed and each of the other cancer types. ORs were estimated per standard deviation increase in the PRS. Models were adjusted for age at specimen collection, first 10 ancestry PCs, sex (except models for sex-specific cancers), reagent kit used for genotyping (Axiom v1 or v2; GERA only), and genotyping array (UK Biobank only). After conducting analyses by cohort, we combined results across cohorts using fixed effects meta-analyses. Heterogeneity was assessed based on *I*^2^ and Cochran’s Q.

For variants contributing to any of the 17 PRS, we estimated the associated risk for each cancer type using logistic regression adjusted for the aforementioned covariables. Variants were modeled individually on a log-additive scale. Results from both cohorts were meta-analyzed. We then visualized the genomic regions that were overrepresented among pleiotropic variants relative to all PRS variants.

In secondary analyses, we used the UK Biobank to explore associations between each PRS and 20 cancer risk factors or serum biomarkers. Logistic (smoking status) or linear (remaining phenotypes) regression models were restricted to cancer-free controls and adjusted for the covariables noted above, as well as cigarette pack-years (forced expiratory volume in 1 second [FEV_1_]/forced vital capacity [FVC]), assay date (serum biomarkers), and use of medications to lower cholesterol (HDL and LDL), control blood pressure (systolic and diastolic blood pressure), and regulate insulin (HbA1c).

All statistical analyses were performed using R 3.2.2 or 3.3.3 (http://www.r-project.org/).

## RESULTS

We abstracted 17,868 genome-wide significant associations from 273 published GWAS (**Supplementary Table 1**). Of the selected set of 880 risk variants independent within the 17 cancer types, 808 variants were independent across all cancer types (**Supplementary Tables 2a-2q**). Endometrial cancer had the fewest independent risk variants (n = 9), and breast cancer had the most (n = 187) (**Figure 1**).

**Figure 1.**
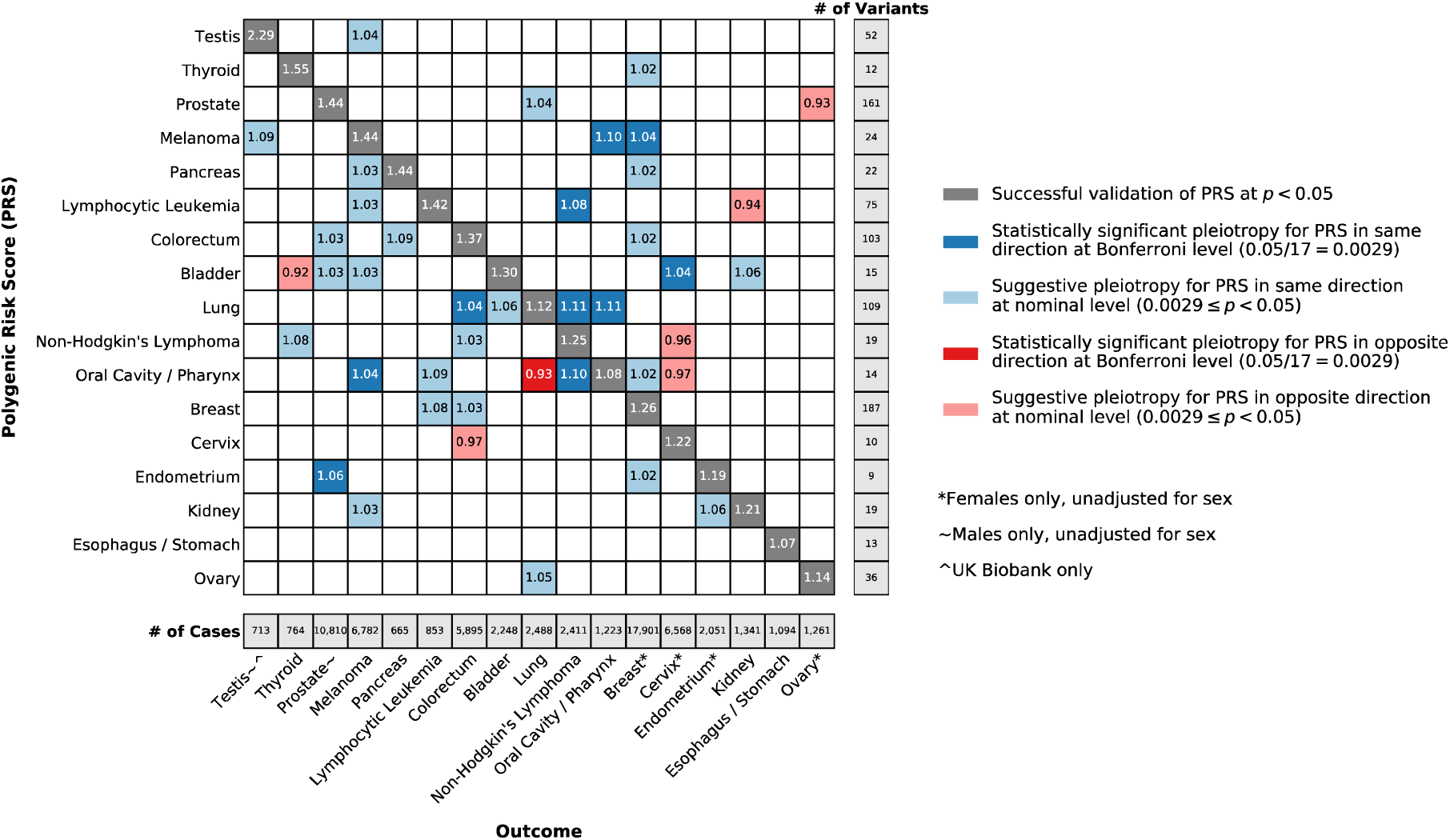
Odds ratios for at least nominally significant associations between cancer-specific polygenic risk scores (PRS) and cancer outcomes, based on meta-analyses of European ancestry participants from the Genetic Epidemiology Research on Adult Health and Aging (GERA) cohort and UK Biobank. Cancers are ordered based on hierarchical clustering of the odds ratios for each PRS across cancer outcomes.

Participants were more commonly female than male (**Supplementary Table 3**). GERA participants were older than UK Biobank participants (mean age: cases, 69 versus 60; controls, 62 versus 57). Case counts ranged from 665 for pancreatic cancer to 17,901 for breast cancer (**Figure 1**). Meta-analyses of non-sex-specific cancers included 410,354 controls. Female-specific meta-analyses included 219,648 controls. Meta-analyses of prostate cancer included 190,706 male controls. For testicular cancer, analyses included 169,967 male controls (UK Biobank only).

Each PRS replicated at a nominal significance level (*p*<0.05; dark gray cells in **Figure 1**) for its corresponding cancer outcome. The largest effect sizes per standard deviation increase in the PRS were observed for testicular (OR=2.29; *p*=6.82×10^−105^) and thyroid cancers (OR=1.55; *p*=6.38×10^−33^). The smallest were observed for esophageal/stomach (OR=1.07; *p*=0.039) and oral cavity/pharyngeal cancers (OR=1.08; *p*=0.007). None of these replicative associations demonstrated significant heterogeneity across cohorts (*p*_Cochran’s-Q_<0.05). **Supplementary Tables 4a, 4b, and 4c** include summary statistics from the meta-analyses, GERA, and UK Biobank, respectively.

Eleven associations between a PRS and cross-cancer outcome were found after correction for multiple testing (*p*<0.05/17=0.0029; **Figure 1**). Results remained materially unchanged correcting for the false discovery rate at *q*<0.05 (**Supplementary Figure 2**). Ten pairs showed a positive association: bladder cancer PRS with cervical cancer (OR=1.04; *p*=9.04×10^−4^); endometrial cancer PRS with prostate cancer (OR=1.06; *p*=5.34×10^−9^); lung cancer PRS with non-Hodgkin’s lymphoma (NHL; OR=1.11; *p*=5.57×10^−7^), colorectal cancer (OR=1.04; *p*=1.22×10^−3^), and oral cavity/pharyngeal cancer (OR=1.11; *p*=1.06×10^−4^); lymphocytic leukemia PRS with NHL (OR=1.08; *p*=1.48×10^−4^); melanoma PRS with breast (OR=1.04; *p*=6.33×10^−7^) and oral cavity/pharyngeal cancers (OR=1.10; *p=*7.84×10^−4^); and oral cavity/pharyngeal cancer PRS with melanoma (OR=1.04; *p*=2.04×10^−3^) and NHL (OR=1.10; *p*=2.67×10^−6^). The oral cavity/pharyngeal cancer PRS was inversely associated with lung cancer (OR*=*0.93; *p*=6.25×10^−4^). Only the melanoma PRS-breast cancer association demonstrated heterogeneity (*I*^*2*^=0.79; *p*_Cochran’s-Q_=0.029). Thirty additional associations (24 positive, six inverse) were nominally significant (*p*<0.05).

Associations between each PRS variant and cancer type are compiled in **Supplementary Tables 5a-5q**. In total, 141 cross-cancer associations were detected at a threshold corrected for the number of effective independent tests (*p*<0.05/808=6.2×10^−5^; **Supplementary Table 6**; includes 18 duplicate associations in which the same variant originated from multiple PRS). They included associations for 55 variants in LD with previously identified risk variants for the outcome cancer. Among the remaining 86 associations, 60 were novel, in that the variant (or variants with *r*^*2*^>0.3 in the 1000 Genomes EUR superpopulation reference panel) had not previously been associated with the outcome cancer at *p*<1×10^−6^ (**Figures 2a and 2b**; includes five duplicate associations originating from multiple PRS). The cancer types with the largest number of novel risk variants were prostate (n = 15), NHL (n = 14; includes one variant originating from multiple PRS), and cervical (n = 12).

**Figure 2.**
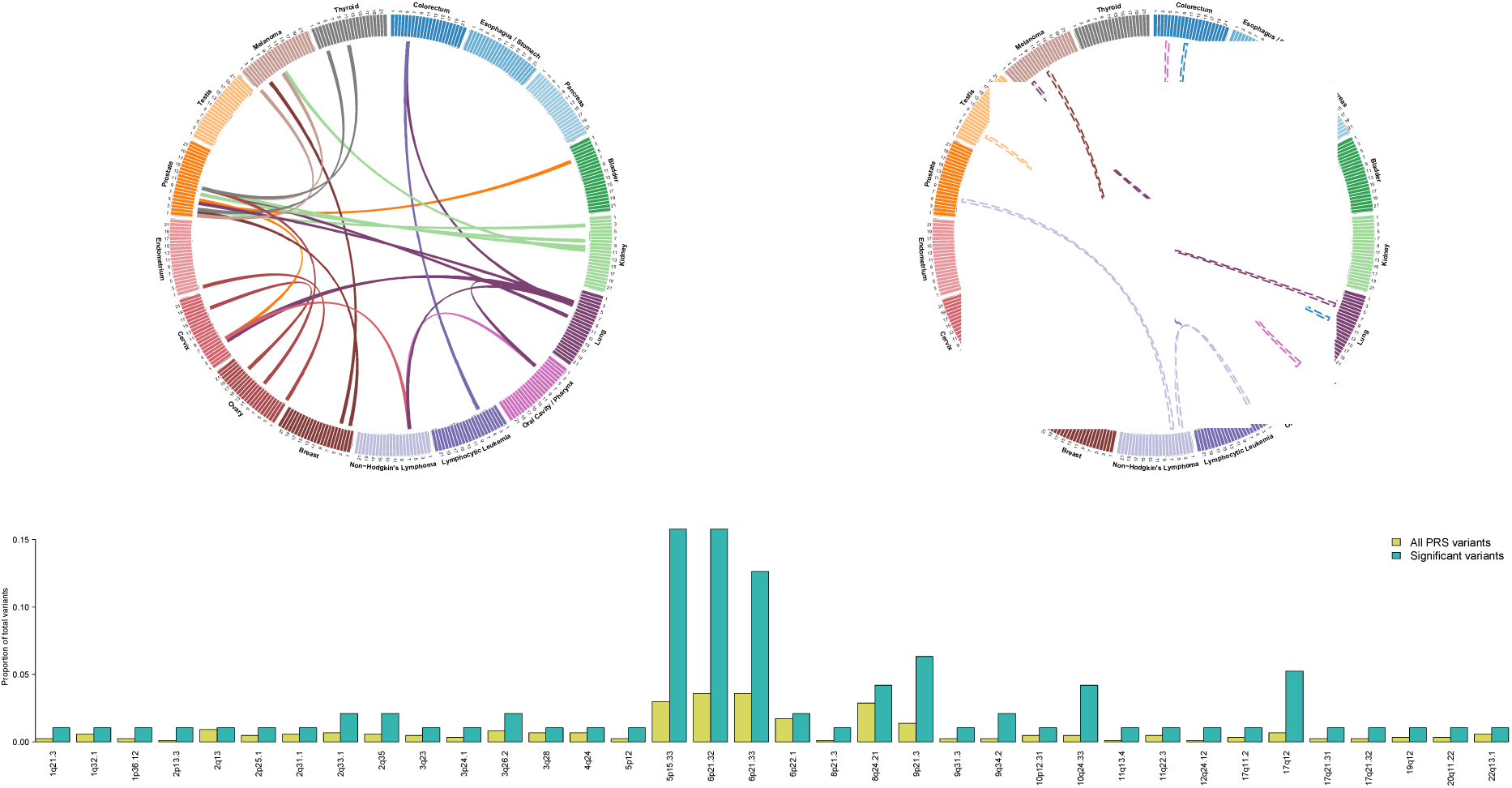
Pleiotropic risk variants from the 17 cancer-specific polygenic risk scores (PRS). (a) Circos plot describing each *positive* association between a known risk variant for one cancer type and a novel cancer phenotype. (b) Circos plot describing each *inverse* association between a known risk variant for one cancer type and a novel cancer phenotype. Each line in (a) and (b) represents a significant association, corrected for the number of effective independent tests (*p*<0.05/808=6.2×10^−5^), between a risk variant for the cancer from which the line originates (denoted by line color) and the cancer type to which the line connects. Cancers are organized by organ site. (c) Region enrichment for 141 significant novel and known associations compared to all PRS variants.

Several genomic regions were overrepresented among pleiotropic variants compared to all PRS variants (**Figure 2c**). Across the 141 cross-cancer associations, pleiotropic variants were most commonly found in *TERT-CLPTM1L* (16% versus 3.0%) and HLA (6p21.32: 16% versus 3.6%; 6p21.33: 13% versus 3.6%). Additional regions enriched for pleiotropy included 9q34.2 (2.1% versus 0.23%), 10q24.33 (4.2% versus 0.46%), 12q24.12 (1.1% versus 0.11%), and 17q12 (5.3% versus 0.69%). These regions remained enriched following normalization by region size (**Supplementary Figure 3**).

Upon evaluating relationships between each cancer PRS and cancer risk factors or biomarkers, we identified 65 statistically significant associations (*p*<0.05/20=0.0025; **Figure 3, Supplementary Table 7**). The lung cancer PRS was associated with the most (12) phenotypes. Positively associated phenotypes included cigarettes per day in smokers (*p*=6.06×10^−32^), pulmonary obstruction (decreasing FEV_1_/FVC; *p*=1.97×10^−25^), HbA1c (*p*=1.59×10^−22^), height (*p*=1.30×10^−4^), and multiple metrics of adiposity (e.g., BMI: *p*=7.63×10^−9^). The lung cancer PRS was associated with lower levels of IGF-1 (*p*=8.58×10^−18^) and HDL cholesterol (*p*=3.94×10^−17^).

**Figure 3.**
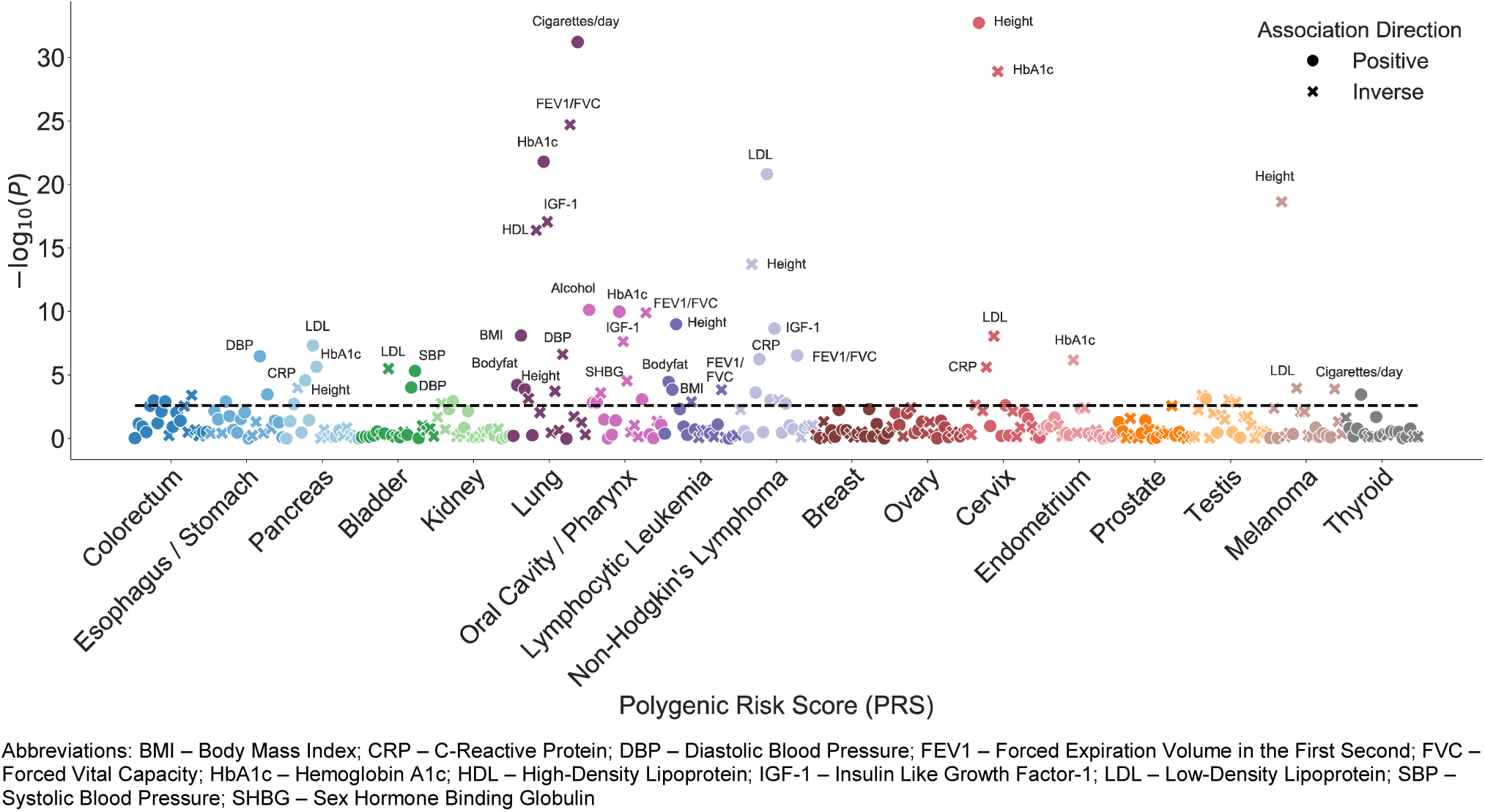
Associations between each cancer-specific polygenic risk score (PRS) and 20 cancer risk factors and related serum biomarkers. All associations were estimated in cancer-free controls in the UK Biobank. Circles denote positive associations between the PRS and the secondary phenotype; crosses denote an inverse direction of association. The dashed line indicates the significance threshold corrected for multiple testing (p<0.05/20 = 0.0025).

The NHL and oral cavity/pharyngeal cancer PRS were each associated with nine secondary phenotypes. Among the associations for the former were increasing levels of LDL (*p*=1.53×10^−21^), IGF-1 (*p*=2.13×10^−9^), and CRP (*p*=5.50×10^−7^). The latter was associated with increasing alcohol intake (*p*=7.28×10^−11^) and pulmonary obstruction (*p*=1.26×10^−10^). PRS for breast, prostate, and ovarian cancers were not clearly associated with any secondary phenotypes. Among the secondary phenotypes, height showed the most cancer PRS associations (n=8; 4 positive, 4 inverse), followed by HbA1c (n=7; 5 positive, 2 inverse), and BMI (4 positive, 2 inverse), pulmonary obstruction (4 positive, 2 inverse), and LDL (2 positive, 4 inverse) (n=6 each).

## DISCUSSION

In this comprehensive study of PRS-specific cancer pleiotropy, we constructed 17 PRS based on systematic review of the cancer GWAS literature. Analyses identified 11 statistically significant cross-cancer PRS associations, as well as novel cancer associations with 55 unique risk variants in known susceptibility regions. We further identified 65 cancer PRS associations with selected non-cancer phenotypes.

Of all PRS evaluated, the oral cavity/pharyngeal and lung cancer PRS were most commonly implicated in associations with cross-cancer and non-cancer phenotypes. These results support existing evidence of cancer pleiotropy, given that the PRS for both cancers included variants in two well-known pleiotropic cancer regions – *TERT*-*CLPTM1L* (2) and HLA (3,4). Notwithstanding shared susceptibility regions, the relationship between oral cavity/pharyngeal and lung cancers was inconsistent. In one direction, the oral cavity/pharyngeal cancer PRS was *inversely* associated with lung cancer. The negative pleiotropy could be partly attributable to two oral cavity/pharyngeal PRS variants (rs467095 and rs10462706; **Supplementary Tables 8 and 9**), both expression quantitative trait loci for *TERT* and *CLPTM1L*, which were inversely associated with lung cancer risk and in LD (*r*^*2*^=0.96 and 0.66, respectively) with variants in the lung cancer PRS. The oral cavity/pharyngeal cancer PRS was also associated with increasing alcohol intake, an established risk factor for such cancers (24). The relationship between alcohol intake and lung cancer remains controversial, with the possibility of an inverse or J-shaped relationship (25,26). In the other direction, the lung cancer PRS was *positively* associated with oral cavity/pharyngeal cancer risk. The positive pleiotropy may be partially explained by the association between the lung cancer PRS and increasing cigarettes per day among smokers. PRS for both cancers were also associated with pulmonary obstruction (i.e., decreasing FEV_1_/FVC), a known lung cancer risk factor (27), as well as higher HbA1c and lower IGF-1 levels, both of which indicate insulin resistance.

Oral cavity/pharyngeal cancer also showed a bidirectional, positive relationship with melanoma, even though the two PRS share only one pair of variants in LD in *TERT*-*CLPTM1L* (**Supplementary Tables 8 and 9**). PRS for both cancer types were inversely associated with height, which is somewhat surprising since increasing height has been strongly associated with melanoma risk (28).

The lung cancer PRS was positively associated with colorectal cancer and NHL. The former association did not appear to be driven by variants in LD; only two out of 109 lung cancer risk variants (rs2853677 and rs1333040) are in high LD (r^2^=0.62 and 0.49, respectively) with colorectal cancer risk variants (rs2735940 and rs1537372, respectively), and neither was strongly associated with colorectal cancer risk in our data. Given that the lung cancer PRS was associated with increasing BMI, body fat, and cigarettes per day, its association with colorectal cancer risk coheres with known risk factors. As five of the lung cancer variants in HLA are in LD with NHL risk variants, LD structure likely played a larger role in the latter association. Both the lung cancer and NHL PRS were associated with increasing HbA1c levels, implicating insulin resistance as a possible shared mechanism. It could be that the NHL PRS was not associated with lung cancer risk because it included only 19 SNPs (relative to 109 SNPs in the lung cancer PRS). We also identified a novel association between a lung cancer risk variant and NHL; rs652888 (6p21.33 in *EHMT2*) has been linked to several autoimmune and infectious diseases (29,30), as well as infection with Epstein-Barr virus (31), a known NHL risk factor (32).

Among the remaining significant cross-cancer PRS associations, two included cancers with PRS variants that were completely independent at the *r*^*2*^=0.3 threshold: the bladder cancer PRS with cervical cancer and the oral cavity/pharyngeal cancer PRS with NHL. Cervical cancer and NHL were among the cancers with the most novel risk variants. Although none of the 15 bladder cancer variants are in LD with known genome-wide significant risk variants for cervical cancer, one *CLPTM1L* variant (rs401681-C) was associated with increased cervical cancer risk at a genome-wide significance level in our study, confirming a suggestive association signal reported previously (33). Similarly, two oral cavity/pharyngeal cancer variants in HLA (rs9271378 and rs3135006), a region that has previously been implicated in NHL (34), were strongly associated with NHL risk in our analyses.

Increasing NHL risk was also associated with the lymphocytic leukemia PRS. Out of 64 lymphocytic leukemia risk variants, only one (rs4987855) is in LD (*r*^*2*^=0.95) with an NHL risk variant (rs17749561). Our results align with those from Sampson, *et al*., which showed an association between a PRS for chronic lymphocytic leukemia (CLL) and the risk of diffuse large B-cell lymphoma, the most common NHL subtype (8). Both CLL and NHL arise from B-cells, and recent classifications account for their similar origin (35).

The association between the endometrial cancer PRS and prostate cancer risk also validated results from Sampson, *et al*. (8) The remaining cross-cancer association from our study – between the melanoma PRS and breast cancer – was not evaluated. Their study did, however, identify two associations that our analyses did not validate: 1) a lung cancer PRS and bladder cancer risk, and 2) an endometrial cancer PRS and testicular cancer risk. Given differences in study design and the many additional SNPs that have been discovered since 2015, it is not especially surprising that some results were distinct.

The genomic regions overrepresented among pleiotropic variants support existing knowledge about shared mechanisms of carcinogenesis. In addition to *TERT-CLPTM1L* and HLA, 9q34.2, 10q24.33, 12q24.12, and 17q12 have been implicated in susceptibility for multiple cancer types. Variants in the breast (36) and pancreatic cancer (37) susceptibility locus 9q34.2 influence estrogen receptor signaling and insulin resistance, and were recently associated with protein biomarkers affecting carcinogenesis (38). The 10q24.33 region containing *OBFC1*, a known telomere maintenance gene, has been implicated in lymphocytic leukemia, melanoma, and kidney, ovarian, and thyroid cancers (39-43). A previous cross-cancer analysis linked 12q24.12 to both colorectal and endometrial cancer risk (44). This locus includes *SH2B3*, a gene involved in regulating signaling pathways related to hematopoiesis, inflammation, and cell migration. The 17q12 locus includes *HNF1B*, which has been extensively characterized with respect to hormonally driven cancers (45).

The non-cancer phenotypes that most frequently surfaced in associations with cancer PRS offer additional mechanistic insights. For example, the lymphocytic leukemia, NHL, and kidney, lung, oral cavity/pharyngeal, and pancreatic cancer PRS were associated with at least one anthropometric trait and showed directionally consistent associations with HbA1c and IGF-1 levels. Obesity-induced chronic inflammation and oxidative stress create a milieu conducive to malignant transformation (46). Furthermore, the metabolic reprogramming necessary to meet the increased energy requirements of proliferating malignant cells is a known hallmark of cancer (1). There is also complex interplay between genetic determinants of adiposity and smoking behaviors (47). Taken together, the findings further implicate obesity-related metabolic dysregulation in cancer susceptibility for multiple sites.

Among the limitations of our study was the inclusion of exclusively European ancestry individuals; results may not be generalizable to diverse populations. We were also limited by modest numbers for some cancers. We favored their inclusion in an effort to evaluate more cancer types than previous investigations. We furthermore combined esophageal and stomach cancers and, separately, oral cavity and pharyngeal cancers into composite phenotypes. While there is precedent for doing so (48-50), we acknowledge the potential resulting phenotypic heterogeneity. We note that our analyses included prevalent and incident cases. However, results from *a posteriori* cross-cancer PRS analyses restricted to incident cases mirrored those from the primary analyses (**Supplementary Table 10**). Our findings are thus unlikely to be driven by associations with survival rather than risk. We also note that our PRS were comprised of exclusively genome-wide significant variants. While a less stringent threshold for inclusion might have yielded more signal, it would not have been based on convincing *a priori* evidence. Finally, while all PRS replicated for their target cancers, some individual risk variants did not. Nevertheless, 92% had effect estimates with consistent directionality relative to the published literature.

Among the strengths of our study was use of two large cohorts with abundant individual-level genetic and phenotypic data, independent of those from which risk variants were identified in prior cancer GWAS (except limited use of UK Biobank data in Huyghe, *et al*.; see Materials and Methods) (20). We also comprehensively reviewed the contemporary literature to identify genome-wide significant risk variants for 17 cancer types. Evaluating risk variants identified for one cancer with respect to risk for others enabled discovery of novel susceptibility loci that would not otherwise meet the strict criteria for genome-wide significance. By additionally evaluating associations with cancer risk factors, we generated insights into pathways that may be influenced by genetic variants implicated in cancer.

Our work expands the repertoire of genetic susceptibility variants for multiple cancers, which should prompt future investigations of their biological and clinical relevance. Although the precise biological mechanisms underpinning the associations remain ambiguous, our findings may still be leveraged toward a more integrated model of cancer risk prediction that considers cross-phenotype effects in addition to cancer-specific risk factors. An approach that incorporates genetic susceptibility profiles may have the greatest potential to aid in risk prediction for cancers with few modifiable risk factors. Combined with future research that investigates pleiotropy in cancer subgroups (e.g., by smoking status or histology) and clinical applications of PRS, our results may inform new strategies toward reducing the burden of cancer.

## ACKNOWLEDGMENTS

Data from the UK Biobank resource was obtained under application number 14105. We thank Alexander Dean for his initial contributions to the development of PRS in GERA.

