## Supplementary Figures 1-3 for "Cross-Cancer Evaluation of Polygenic Risk Scores for 17 Cancer Types in Two Large Cohorts"

**Supplementary Figure 1.** Flow chart depicting the reduction of genome-wide significant associations abstracted from the literature to the variants included in the polygenic risk scores (PRS)

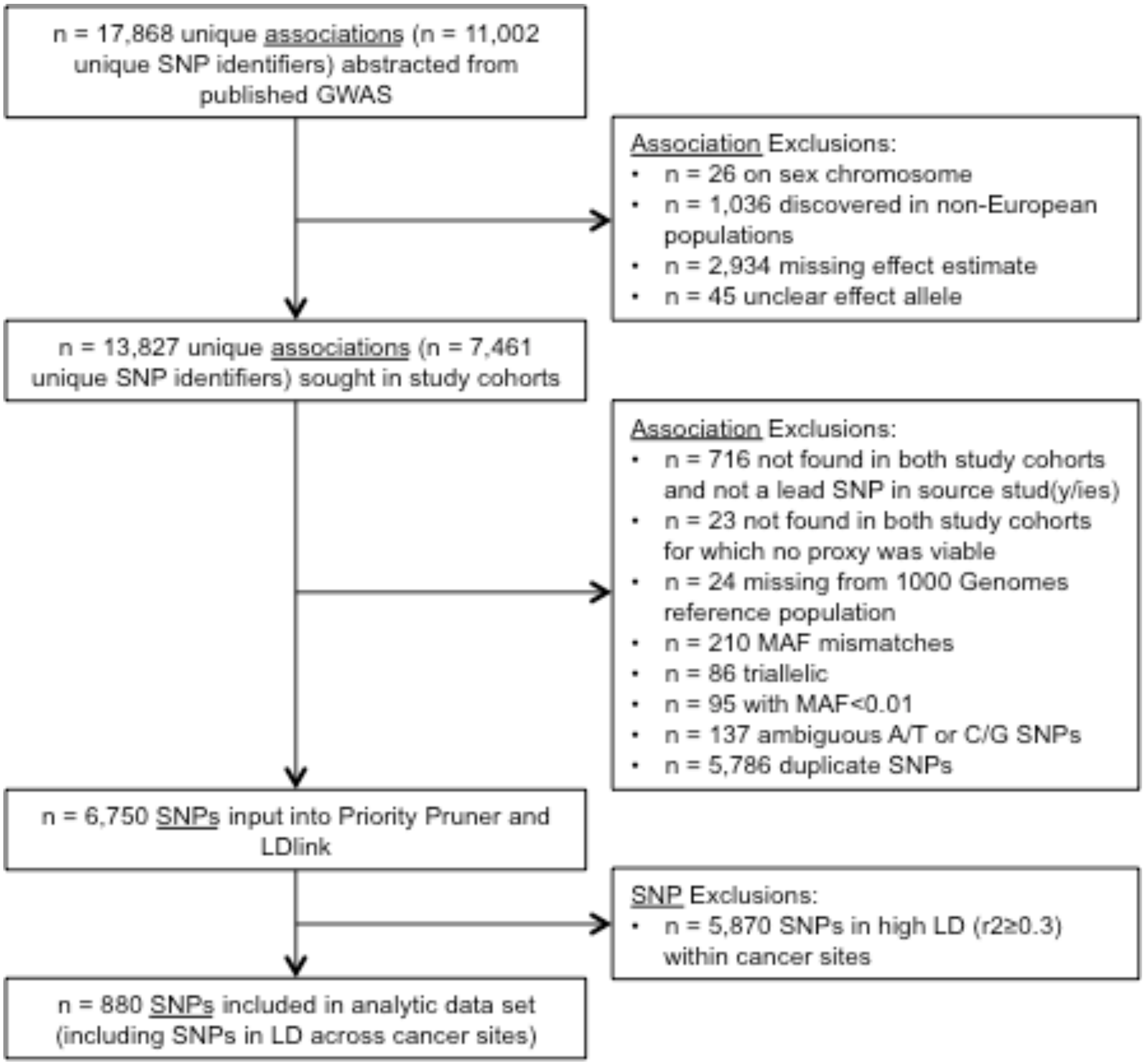

**Supplementary Figure 2.** Odds ratios for significant associations between cancer-specific polygenic risk scores (PRS) and cancer outcomes, based on meta-analyses of European ancestry participants from the Genetic Epidemiology Research on Aging (GERA) cohort and UK Biobank, according to a false discovery rate threshold of *q*<0.05. Cancers are ordered based on clustering of the odds ratios for each PRS across cancer outcomes.

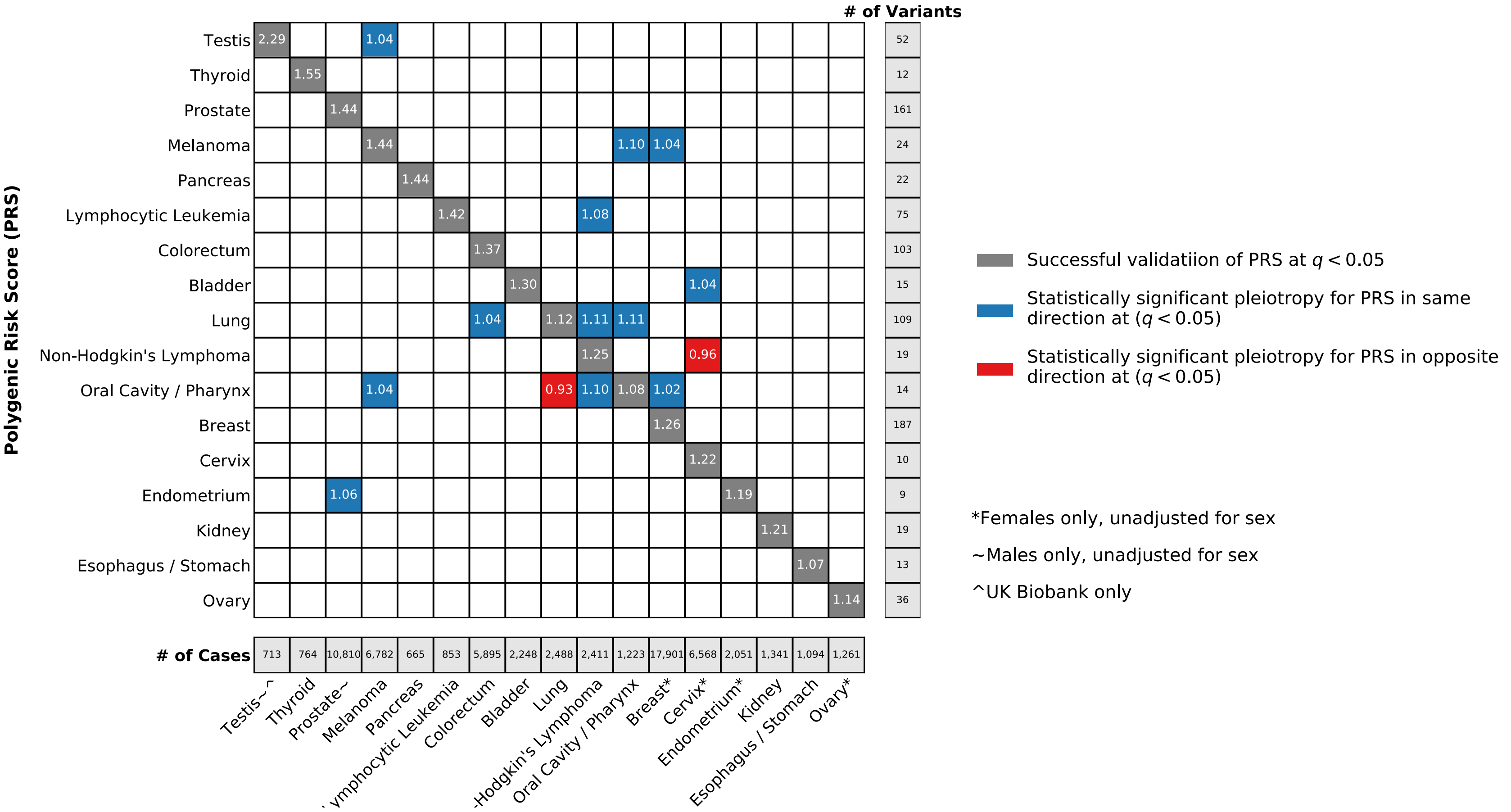

**Supplementary Figure 3.** Region enrichment for 141 significant novel and known pleiotropic risk variants compared to all PRS variants. The proportion of significant variants in each region is normalized by the region size relative to the genome size per kb.

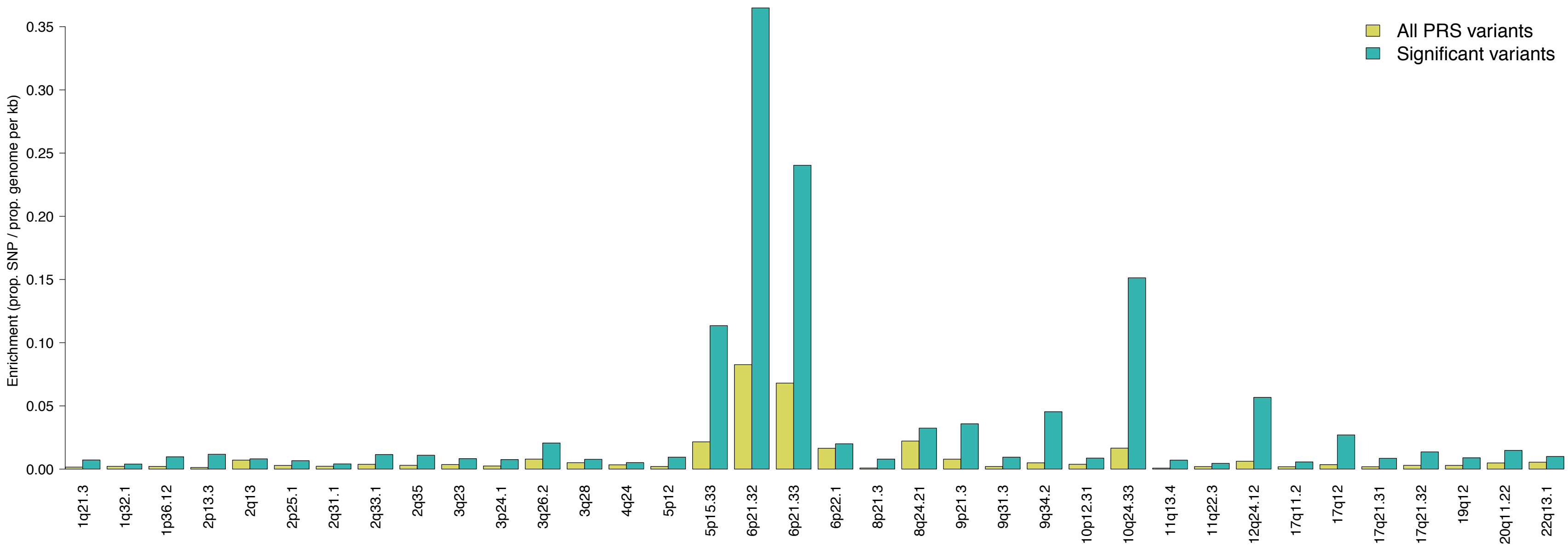
